# Infection of brain pericytes underlying neuropathology of COVID-19 patients

**DOI:** 10.1101/2021.05.24.445532

**Authors:** Matteo Bocci, Clara Oudenaarden, Xavier Sàenz-Sardà, Joel Simrén, Arvid Edén, Jonas Sjölund, Christina Möller, Magnus Gisslén, Henrik Zetterberg, Elisabet Englund, Kristian Pietras

## Abstract

A wide range of neurological manifestations have been associated with the development of COVID-19 following SARS-CoV-2 infection. However, the etiology of the neurological symptomatology is still largely unexplored.

Here, we used state-of-the-art multiplexed immunostaining of human brains (*n* = 6 COVID-19, median age = 69,5 years; and *n* = 7 control, median age = 68 years), and demonstrated that expression of the SARS-CoV-2 receptor ACE2 is restricted to a subset of neurovascular pericytes. Strikingly, neurological symptoms were exclusive to, and ubiquitous in, patients that exhibited moderate to high ACE2 expression in peri-vascular cells. Viral particles were identified in the vascular wall and paralleled by peri-vascular inflammation, as signified by T cell and macrophage infiltration. Furthermore, fibrinogen leakage indicated compromised integrity of the blood-brain barrier. Notably, cerebrospinal fluid from an additional 16 individuals (*n* = 8 COVID-19, median age = 67 years; and *n* = 8 control, median age = 69,5 years) exhibited significantly lower levels of the pericyte marker PDGFRβ in SARS-CoV-2-infected cases, indicative of disrupted pericyte homeostasis.

We conclude that pericyte infection by SARS-CoV-2 underlies virus entry into the privileged central nervous system space, as well as neurological symptomatology due to peri-vascular inflammation and a locally compromised blood-brain barrier.

## Introduction

The clinical manifestations of coronavirus disease 2019 (COVID-19) infection primarily include respiratory symptoms, ranging from a mild cough to severe bilateral pneumonia^1, 2^. However, SARS-CoV-2 bears an organotropism beyond the respiratory tract^3, 4^, with increasing testimony indicating the brain as an extrapulmonary target of SARS-CoV-2^5^. The involvement of the CNS encompasses a broad spectrum of neurological manifestations (including headache, fatigue, anosmia, ageusia, confusion, loss of consciousness), often representing an ulterior clinical morbidity that significantly contributes to COVID-19-related deaths^6–8^.

The main entry receptor for SARS-CoV-2 is reported to be the angiotensin-converting enzyme (ACE)2, which is a component of the renin-angiotensin system^9, 10^. To date there is still no conclusive evidence concerning the localization of ACE2 in the human CNS^3^, and the mechanism of SARS-CoV-2 infection in the brain remains a conundrum.

Here, using highly sensitive multiplexed immunohistochemistry (mIHC) of brain tissue from a series of confirmed COVID-19 patients and corresponding controls, we determined that ACE2 is exclusively expressed by brain pericytes in the subset of patients that also exhibited neurological symptoms. Moreover, spatial immunophenotyping revealed a localized perivascular inflammation in brain tissue from COVID-19 patients, paralleled by an impairment of the functionality of the vascular wall as indicated by loss of integrity of the blood-brain barrier (BBB). Finally, in the cerebrospinal fluid (CSF) of a cohort of COVID-19 patients with neurological involvement, levels of soluble PDGFRβ, a pericyte-specific marker in the brain, were significantly reduced compared with non-COVID-19 individuals, suggestive of SARS-CoV-2-related functional impairment of pericytes. Taken together, our findings highlight a previously unappreciated role for brain pericytes in acting as pioneers for SARS-CoV-2 entry into the CNS.

## Materials and methods

### Patients

Excessive brain tissues sampled from six COVID-19 autopsies and 7 non-COVID-19 cases were used to create formalin-fixed paraffin-embedded (FFPE) blocks (Supplementary Table 1). The use of these samples was approved by the Central Ethical Review Authority in Sweden (2020-02369, 2020-06582, and 2020-01771). Clinical data with details of neurologic symptoms or other signs of brain affection were sought in the referral documents or else in the Regional Medical Records database Melior, which was used also for the diagnostic work-up.

CSF from eight patients with neurological manifestations admitted to the Sahlgrenska University Hospital in Gothenburg, Sweden was included (Supplementary Table 2). Infection with SARS-CoV-2 was confirmed *via* rtPCR analysis. Age- and sex-matched non-COVID-19 controls were selected, consisting of patients who were examined because of clinical suspicion of neurological disease, but where no neurochemical evidence was found, based on clinical reference intervals. The use of these samples has been approved by the Regional Ethical Committee in Gothenburg.

### Bioinformatics data access and analysis

Expression of *Ace2* was investigated in publicly available single cell RNA sequencing (scRNA-seq) SMART-Seq2 RNA-seq libraries on FACS-sorted non-myeloid brain cells of seven mice (*Tabula Muris*)^11^, and in a database of murine vasculature^12, 13^.

Mouse whole brain and hippocampus SMART-seq data (gene expression aggregated per cluster, calculated as trimmed means) from the Allen Brain Atlas consortium was downloaded on 14^th^ October 2020^14, 15^. For expression of Pvalb and Sst neurons, the average was calculated of 13 and 40 cell clusters, respectively.

Human ACE2 protein expression images were retrieved from the Human Protein Atlas initiative (Version 20.0)^16^.

### Immunohistochemistry (IHC)

Five μm-thick FFPE tissue sections were dewaxed and rehydrated through xylene and water-based ethanol solutions. Heat-induced epitope retrieval was performed with a pressure cooker (2100 Antigen Retriever, BioVendor, Germany) in citrate or Tris-EDTA buffer (Agilent Dako, USA). Following endogenous peroxidase quenching (BLOXALL, Vector Laboratories, USA), tissues were incubated with CAS-block (Thermo Fisher Scientific, USA), 1 hour at room temperature (RT), and Ultra V block (Thermo Fisher Scientific, USA) for 5 minutes. Primary antibodies (Supplementary Table 3) diluted in CAS-block were applied for 30 minutes, followed by UltraVision ONE HRP polymer (Thermo Fisher Scientific, USA), 30 minutes, at RT. The ImmPACT DAB substrate (Vector Laboratories, USA) was applied. Tissues were counterstained with hematoxylin, dehydrated and mounted with Cytoseal 60 (Thermo Fisher Scientific, USA). Imaging was performed with an automated BX63 microscope connected to a DP-80 camera (Olympus, Japan).

### Multiplexed IHC (mIHC)

FFPE sections used for IHC were subjected to multiplexed labelling following optimized protocols established in the lab. All reagents are from Akoya Biosciences (USA), including the Vectra Polaris scanner for imaging and the PhenoChart/InForm software. Following slide preparation, sections underwent staining cycles (Supplementary Table 4) —including blocking, primary antibody incubation, HRP tagging and labeling with OPAL-conjugated tyramide substrate— and a stripping procedure to remove unbound primary antibody/HRP. A counterstain with DAPI preceded the mounting with ProLong Diamond antifade (Thermo Fisher Scientific, USA).

The composite images were generated by removing inherent autofluorescence signal from an unstained section, as well as by comparing fluorescence intensities to those of a spectral library.

### Soluble PDGFRβ ELISA

CSF sPDGFRβ concentration was measured by sandwich ELISA (Thermo Fisher Scientific, USA), as previously described^17^. Statistical Mann-Whitney U-test was performed using Prism (GraphPad Software, USA). The significance level was set at *P* < 0.05, two-sided.

### Data availability

The authors confirm that the data supporting the findings of this study are available within the article and its supplementary material.

## Results

### The ACE2 receptor is expressed by pericytes in murine and human brain

Expression of ACE2 in the brain has variably been reported in neurons, glial cells, and vascular cells^18–22^. Because of this ambiguity of localization, we started by exploring ACE2 expression in publicly available mRNA and protein datasets from murine and human brain. Mining of the Allen Mouse Brain Atlas of single cell transcriptomes demonstrated unique enrichment for *Ace2* transcript in pericytes (Fig. 1A). A similar compartmentalization was observed in the *Tabula Muris*^11^ as well as in an scRNA-seq compendium of the murine brain vasculature^12, 13^ (Supplementary Fig. 1A and B). In agreement with the transcriptional data, localization of the ACE2 protein by the Human Protein Atlas^16^ was restricted to the peri-vascular compartment in a subset of blood vessels in human cerebral cortex (Supplementary Fig. 1C).

**Figure 1.**
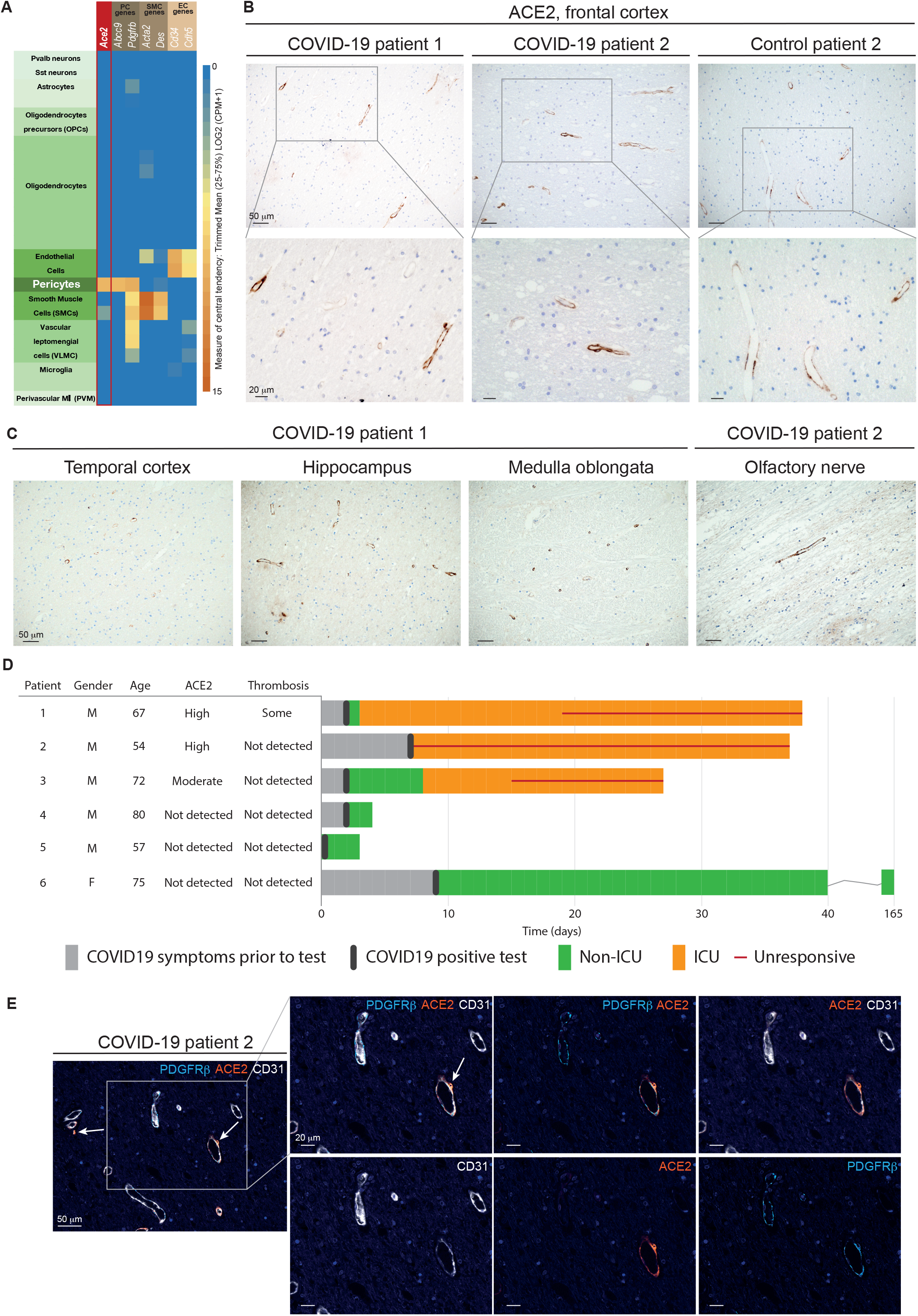
The ACE2 receptor is expressed by pericytes in murine and human brain. (**A**) Expression of *Ace2* in then annotated cell types in the mouse brain based on the Allen Mouse Brain Atlas. (**B**) Representative IHC staining of peri-vascular ACE2 in the frontal cortex of two COVID-19 patients and one control individual. Cell nuclei are counterstained with hematoxylin (blue). (**C**) Representative IHC staining of peri-vascular ACE2 in different brain regions of two COVID-19 patients. Cell nuclei are counterstained with hematoxylin (blue). (**D**) Clinical course of the COVID-19 patients included in the study. For each patient, appearance of symptoms, hospitalization, infection status and progression to death are included, together with the post-mortem evaluation of ACE2 immunoreactivity and thrombosis in the CNS. (**E**) Four-plex mIHC staining of the frontal cortex of a COVID-19 patient. The composite image depicts CD31 (endothelial cells, white), PDGFRβ (pericytes, cyan), and ACE2 (orange). Cell nuclei are counterstained with DAPI (blue). In the inlets, the white arrows indicate ACE2-positive signal in the abluminal side of CD31. The intensity of each individual OPAL fluorophore, and the combined PDGFRβ/ACE2 and CD31/ACE2 overlays are presented.

### The ACE2 protein is expressed by peri-vascular cells of neural tissue from COVID-19 patients with neurological symptoms

Next, we sought to investigate the expression of ACE2 in brain tissue of COVID-19 patients. To this end, we obtained FFPE samples of multiple brain regions from six patients whose death was confirmed to be a consequence of SARS-CoV-2 infection (Supplementary Table 1). In the frontal cortex, moderate to high ACE2 immunoreactivity revealed a vascular pattern in a subset of blood vessels in five of the 13 cases (Fig. 1B). Reassuringly, other brain regions showed equivalent distribution of ACE2, indicating that ACE2 was widely expressed in peri-vascular cells throughout the CNS (Fig. 1C). Notably, ACE2 reactivity, which was confirmed with two different antibodies in positive control tissues from kidney (Supplementary Fig. 1D), appeared to be a patient-specific feature, since some cases did not show signal at all, or with very low frequency (Fig. 1D and Supplementary Fig. 1E). To conclusively validate which cell type harbored ACE2 expression, we performed mIHC on human brain tissue to simultaneously visualize ACE2, CD31^+^ endothelial cells, PDGFRβ^+^ pericytes. ACE2 expression coincided with that of PDGFRβ, but not with CD31 staining (Fig. 1E and Supplementary Fig. 1F). Pericytes investing the vasculature exhibited a nuanced pattern of PDGFRβ and ACE2 immunoreactivity, with some cells bearing positivity solely for PDGFRβ, while others expressed both markers. Remarkably, the three COVID-19 patients that exhibited moderate to high peri-vascular ACE2 expression in the brain all presented with neurological symptoms, while all ACE2-negative patients remained free from such manifestations (Fig. 1D). Collectively, our data demonstrate that in the brain, ACE2 is exclusively expressed by pericytes in a manner that signifies development of neurological symptoms from COVID-19.

### SARS-CoV-2 is detectable in human brain of COVID-19 patients

An increasing body of evidence converges on the inherent difficulty of detecting SARS-CoV-2 in the brain^23, 24^. To build on previous reports on the localization of SARS-CoV-2 in human brain tissue, we additionally analyzed brain samples from non-infected individuals to enable conclusions about the presence of the spike protein or the nucleocapsid protein of SARS-CoV/SARS-CoV-2 in the CNS with a higher certainty. For both viral components, positive areas in brain sections of COVID-19 patients exhibited comparable patterns with those shown in previous studies.^25^ Notably, however, we demonstrated an analogous intensity and distribution of the viral proteins when we probed brain tissues from non-infected individuals (Fig. 2A). In order to unequivocally define our ability to visualize viral particles in human tissues, we gained access to placental tissue from a confirmed case of SARS-CoV-2 vertical transmission to serve as a positive control^26^. We also made use of the J2 antibody specifically designed to detect viral double-stranded (ds)RNA. In the placenta, a 7-plex mIHC panel confirmed the epithelial cytokeratin^+^ syncytiotrophoblasts as the main target for viral infection by virtue of expression of ACE2 and the presence of dsRNA in a well-defined dotted pattern (Fig. 2B and Supplementary Fig. 2A); a pattern of distribution which was essentially preserved with antibodies against the coronaviridae family or SARS-CoV-2-specific antigens (Supplementary Fig. 2B). Finally, applying the now validated protocol for detection of viral dsRNA to brain sections, we identified an analogous dotted pattern in discrete peri-vascular, non-endothelial, cells in the brain of COVID-19 patients (Fig. 2C and Supplementary Fig. 2C). Reassuringly, the peri-vascular staining pattern was absent from brain samples of non-infected individuals. Together with our observations of ACE2 expression in pericytes, our conclusive localization of viral dsRNA suggests that brain pericytes are indeed uniquely susceptible to viral infection and may serve as CNS entry points for SARS-CoV-2.

**Figure 2.**
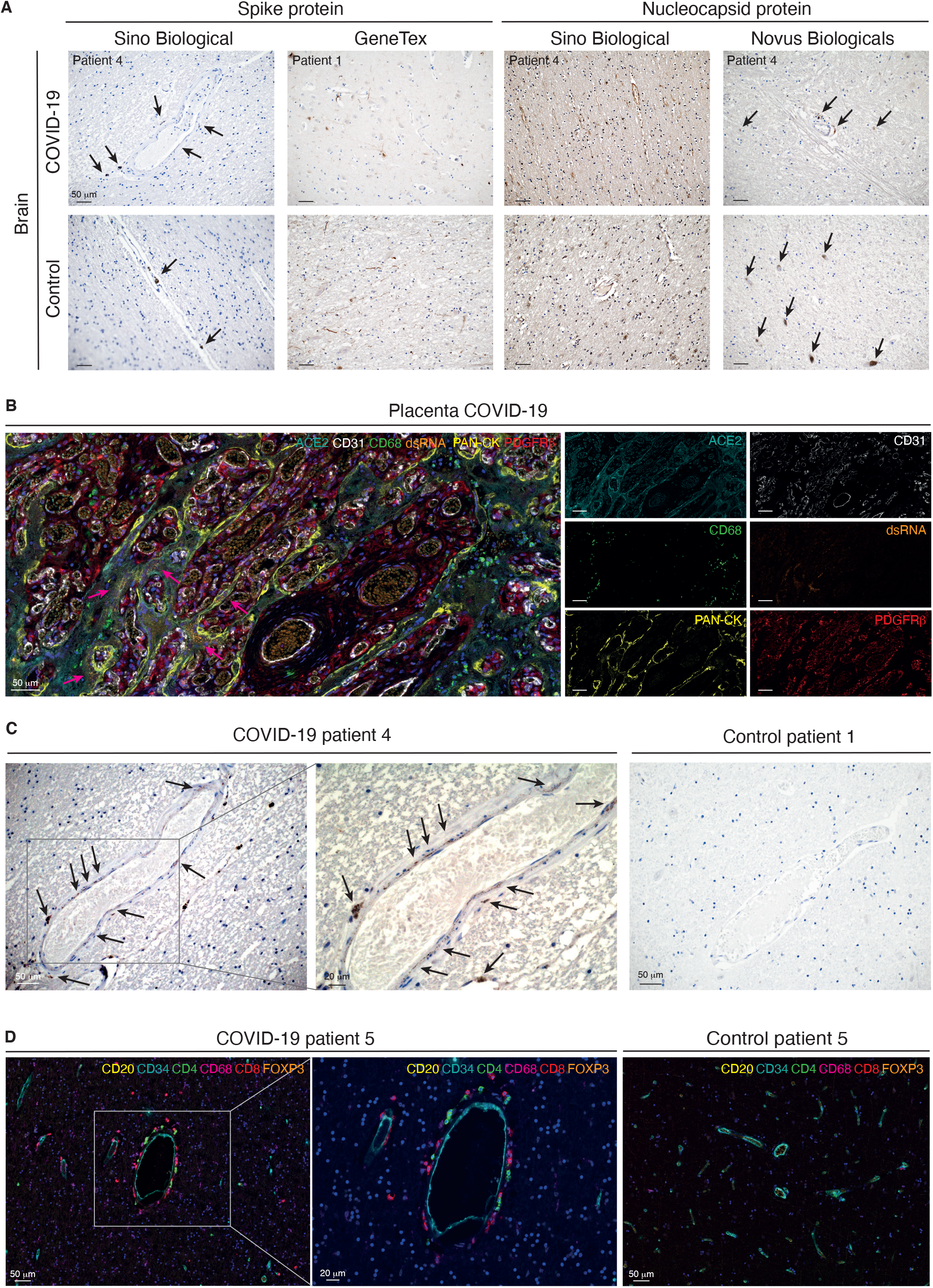
Peri-vascular infection by SARS-CoV-2 is paralleled by peri-vascular inflammation in the brain of COVID-19 patients. (**A**) Immunohistochemical detection of viral components in COVID-19-infected patients and non-COVID-19 controls. Cell nuclei are counterstained with hematoxylin (blue). The black arrows indicate the chromogenic deposition of the 3,3’-diaminobenzidine (DAB) substrate. (**B**) Representative field of a 7-plex mIHC staining panel of placental tissue infected with SARS-CoV-2. The magenta arrows indicate accumulation of viral dsRNA in correspondence of the ACE2-positive areas by the specialized epithelial layer of syncytiotrophoblast in the placenta. The intensity of each OPAL fluorophore is further presented in individual photomicrographs. (**C**) Immunohistochemical detection of dsRNA in the cerebral cortex of a COVID-19 patient and in a non-COVID-19 control. Cell nuclei are counterstained with hematoxylin (blue). Black arrows indicate deposition of the DAB substrate. (D) Composite mIHC image of the peri-vascular immune cell infiltration in the frontal cortex of a COVID-19 patient and in a control individual. The antibody panel was designed for the concomitant detection of CD34 (endothelium) and five immune cell markers: CD4 (T helper cells), CD8 (cytotoxic T lymphocytes), CD20 (B cells), CD68 (macrophages), and FOXP3 (regulatory T cells).

### Peri-vascular infection by SARS-CoV-2 in the brain is paralleled by perivascular inflammation

We hypothesized that infection of pericytes would result in neuro-inflammation and therefore implemented a spatial immunophenotyping approach for the concomitant detection of the endothelium (CD34^+^) and five immune cell populations, including T helper and cytotoxic T lymphocytes, regulatory T cells, B cells, and macrophages. Surrounding the brain vasculature in COVID-19 patients, we detected CD4^+^ and CD8^+^ T cells, as well as CD68^+^ macrophages, indicative of peri-vascular inflammation, rather than widespread neuroinflammation in the brain parenchyme (Fig. 2D and Supplementary Fig. 2D). The immune infiltration did not affect all blood vessels, indicating that the inflammation was not the result of systemic mediators, but rather of local instigation.

### Pericyte infection leads to vascular fibrinogen leakage in the CNS

Next, we investigated whether impaired pericyte function consequential to SARS-CoV-2 infection and the peri-vascular inflammation impinged on the integrity of the vascular wall. We first performed a 7-plex mIHC staining focusing on the permeability of the neurovascular unit. Remarkably, in COVID-19 patients, extra-vascular fibrinogen was readily detected as a characteristic gradient in subsets of vessels, occasionally also characterized by ACE2 expression and the presence of viral dsRNA (Fig. 3A and Supplementary Fig. 3 A and B). Conversely, fibrinogen was fully retained within the blood vessels of non-infected control cases. Moreover, astrocyte priming indicative of local activation of the brain parenchyme was not apparent during COVID-19 infection (Fig. 3B and Supplementary Fig. 3C). Together with our identification of SARS-CoV-2 and immune cell infiltrates in the peri-vascular region, the leakage of fibrinogen from the blood vessels strongly suggests that viral infection of pericytes breaches the tightly organized BBB.

**Figure 3.**
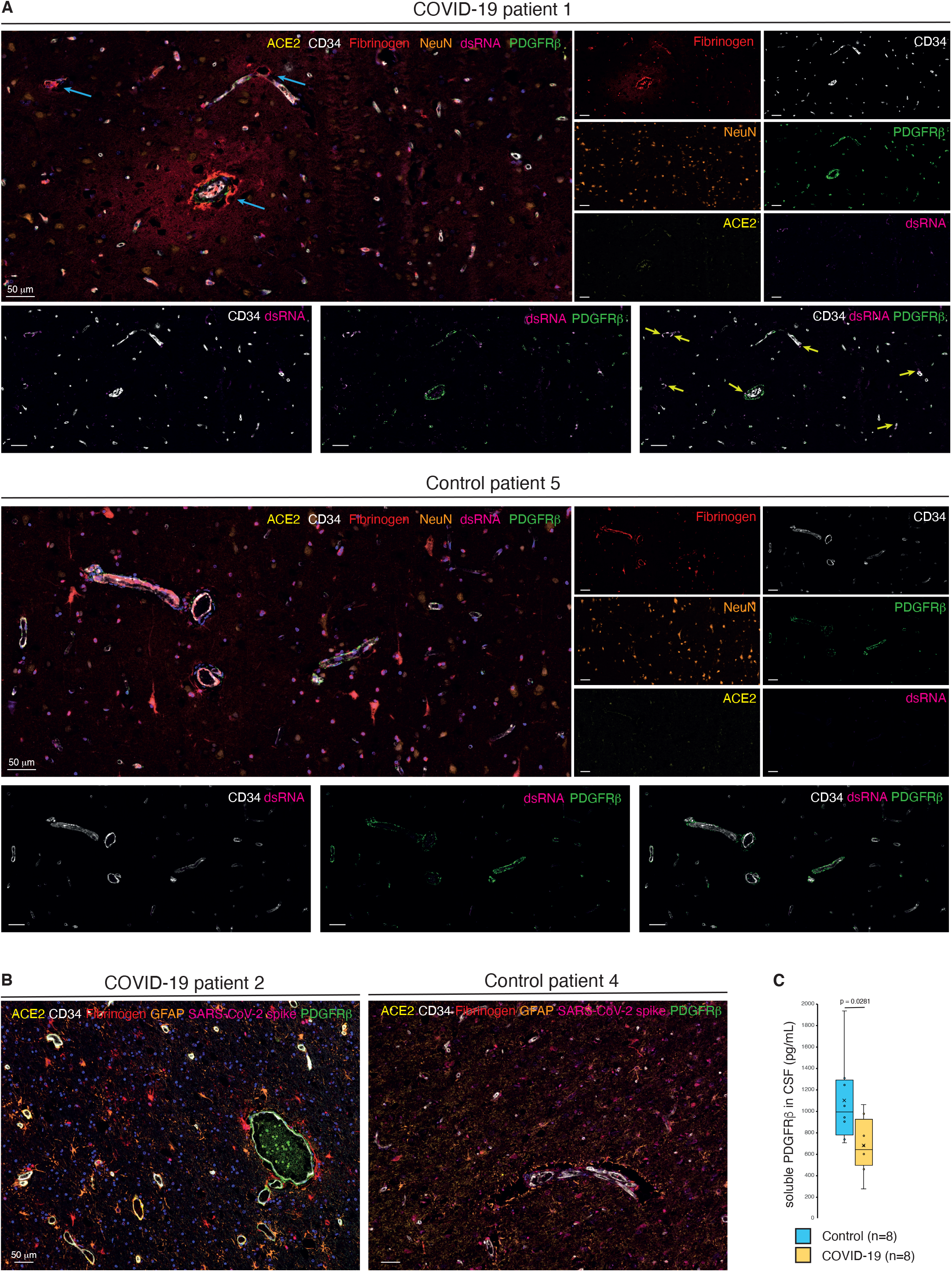
Pericyte infection leads to vascular leakage in the CNS. (**A**) Composite mIHC of the frontal cortex of a COVID-19 patient and a control individual. The fields highlight the fibrinogen halo surrounding leaky blood vessels following SARS-CoV-2 infection. The images depict the neurovascular unit (CD34, PDGFRβ, and ACE2), fibrinogen, viral dsRNA, and neurons. The intensity of each OPAL fluorophore is further presented in individual photomicrographs. The cyan arrows indicate fibrinogen leakage, yellow arrows highlight points of converging PDGFRβ/dsRNA staining. (**B**) Composite mIHC of the frontal cortex of a COVID-19 patient and a control individual. The fields focus on astrocytes priming as a readout of local neuroinflammation. The images depict the neurovascular unit (CD34, PDGFRβ, and ACE2), fibrinogen, SARS-CoV-2 spike protein, and astrocytes. (**C**) boxplot of the concentration of soluble PDGFRβ (pg/ml) in the CSF of COVID-19 patients and non-COVID-19 controls (circles: individual measurements, cross: cohort average).

### Shedding of PDGFRβ into the CSF is reduced in COVID-19 patients

Our findings led us to speculate that the homeostatic state of brain pericytes would be disrupted in COVID-19 patients. Therefore, we collected CSF from an additional eight patients with acute COVID-19 that presented with neurological manifestations, as well as non-infected matched controls (Supplementary Table 2). Intriguingly, the level of the pericyte marker sPDGFRβ in the CSF of COVID-19 patients was on average significantly lower than in non-COVID-19 control individuals as measured by ELISA, indicative of a perturbed pericyte homeostasis (Fig. 3C).

## Discussion

The primary cellular receptor for SARS-CoV-2 entry is ACE2^9^, but the expression pattern of ACE2 in the CNS has not been conclusively resolved. Notably, the few published studies detailing the expression of ACE2 and/or SARS-CoV-2 protein in the CNS lack reliable and appropriate controls, precluding firm conclusions. Here, by means of highly sensitive mIHC and the use of both positive and negative control tissues, we were able to confirm that ACE2 exhibited an exclusive peri-vascular expression pattern in the CNS. Similarly, viral particles and their dsRNA were observed in CNS pericytes in COVID-19 patients, independently of the peri-vascular ACE2 expression status. Whether other co-receptors for SARS-CoV-2, including TMPRSS2, CD147, and neuropilin-1, contribute to CNS-tropism remains to be investigated.

Based on our observations, we hypothesize that infection and subsequent damage of brain vascular pericytes by SARS-CoV-2 and peri-vascular inflammation may lead to impairment of the BBB, instigating neurological complications and possibly virus entry into the CNS. In line with our report, two recent studies observed vascular leakage and peri-vascular immune infiltration in the brain of COVID-19 patients, but without the crucial link to ACE2 expression by, and infection of, pericytes^27, 28^. However, it is still an outstanding question whether SARS-CoV-2 is overtly neurotropic or if the neurological symptoms associated with COVID-19 are secondary to events related to the systemic host response^29^. Although solely based on the comparable abundance of GFAP in the tissues, our observations do not provide support for the hypothesis of a cytokine storm. Nevertheless, immune activation markers β2-microglobulin and neopterin were previously found to be elevated in the CSF of COVID-19 patients^30^. Hence, further investigations are warranted to fully clarify whether a systemic inflammatory response is associated with neurological manifestations of COVID-19.

Intriguingly, COVID-19 patients with neurological symptoms presented with a reduced concentration of pericyte-derived sPDGFRβ in the CSF. While our mIHC of brain tissue demonstrated a surprisingly variable occurrence of PDGFRβ^+^ peri-vascular cells, in line with the results from the CSF analysis, the analysis did not support an overall diminished pericyte coverage of the vasculature of COVID-19 patients. A second, and perhaps more likely, explanation for the reduced expression/shedding of PDGFRβ in COVID-19 patients, is that SARS-CoV-2 infection of pericytes diverted the protein synthesis machinery to produce viral proteins, leading to loss of endogenous marker expression^31^ and consequential functional impairment.

An improved understanding of SARS-CoV-2 neurotropism is urgently needed to guide the clinical management of acute neurological symptoms, as well as to define strategies to prevent post-infectious neurological complications. We propose that a possible entry site of SARS-CoV-2 into the CNS goes through ACE2-expressing pericytes. Interestingly, although overt endothelial cell infection by SARS-CoV-2 does not appear to occur ^22^, a recent investigation determined that radiolabeled S1 spike viral protein could be retained on the abluminal side of endothelial cells where it associated with the capillary glycocalyx in mice or further sequestered by the endothelium^32^. It is thus tempting to speculate that this represents one plausible way to expose pericytes to the viral infection. Furthermore, the absence of brain pericytes in mice results in a disrupted BBB associated with wide-spread loss of integrity^33^. Conversely, sealing of the BBB following thrombolysis after ischemic stroke has been achieved in clinical trials by treatment with the tyrosine kinase inhibitor imatinib^34, 35^. Whether similar interventions aiming to support the integrity of the BBB would alleviate neurological symptoms in COVID-19 patients warrants further studies.

## Abbreviations

BBB: blood-brain barrier
COVID-19: coronavirus disease 2019
CSF: cerebrospinal fluid
HIER: heat-induced epitope retrieval
mIHC: multiplexed immunohistochemistry
SARS-CoV-2: severe acute respiratory syndrome coronavirus 2

## Acknowledgements

We gratefully acknowledge Professor David Gisselsson Nord (Lund University) for provision of placenta tissue specimens, and associate professor Emma Hammarlund (Lund University) for critically reading the manuscript.

## Supplementary material

Supplementary material is available at *Brain* online.

---

**Supplementary Figure 1.**
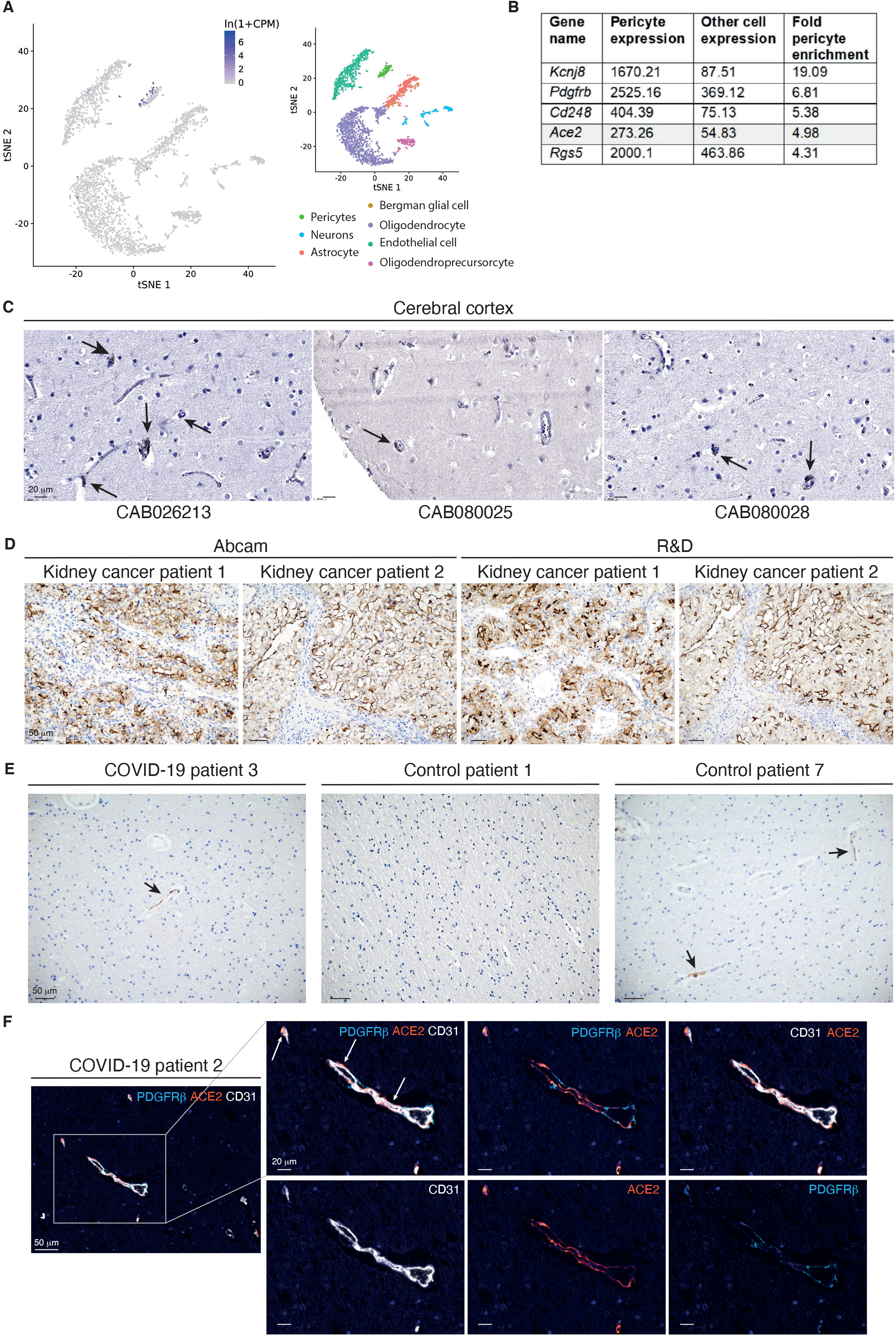
(**A**) Feature plot of *Ace2* expression in seven distinct color-coded cell ontology classes (tissue reference: brain non-myeloid) from the *Tabula muris* compendium. (**B**) Enrichment for *Ace2* in mouse brain pericytes compared with other brain cell types based on the Betsholtz Atlas. (**C**) representative IHC staining of ACE2 in the cerebral cortex of three patients (and three antibodies) from the Human Protein Atlas initiative. Cell nuclei are counterstained with hematoxylin (blue). The black arrows indicate ACE2+ areas. (**D**) Representative IHC staining of ACE2 with two distinct antibodies in two different biopsies of human renal cell carcinoma. Cell nuclei are counterstained with hematoxylin (blue). (**E**) Representative IHC staining of peri-vascular ACE2 in the frontal cortex of one COVID-19 patients and two control individuals. Cell nuclei are counterstained with hematoxylin (blue). The black arrows indicate the chromogenic deposition of the DAB substrate. (**F**) Additional field of the 4-plex mIHC staining of the frontal cortex of the COVID-19 patient in Figure 1E. The final composite image depicts CD31 (endothelial cells, white), PDGFRβ (pericytes, cyan), and ACE2 (orange). Cell nuclei are counterstained with DAPI (blue). In the inlets, the white arrows indicate ACE-positive signal in the abluminal side of CD31. The intensity of each individual OPAL fluorophore, and the combined PDGFRβ/ACE2 and CD31/ACE2 overlays are presented.

**Supplementary Figure 2.**
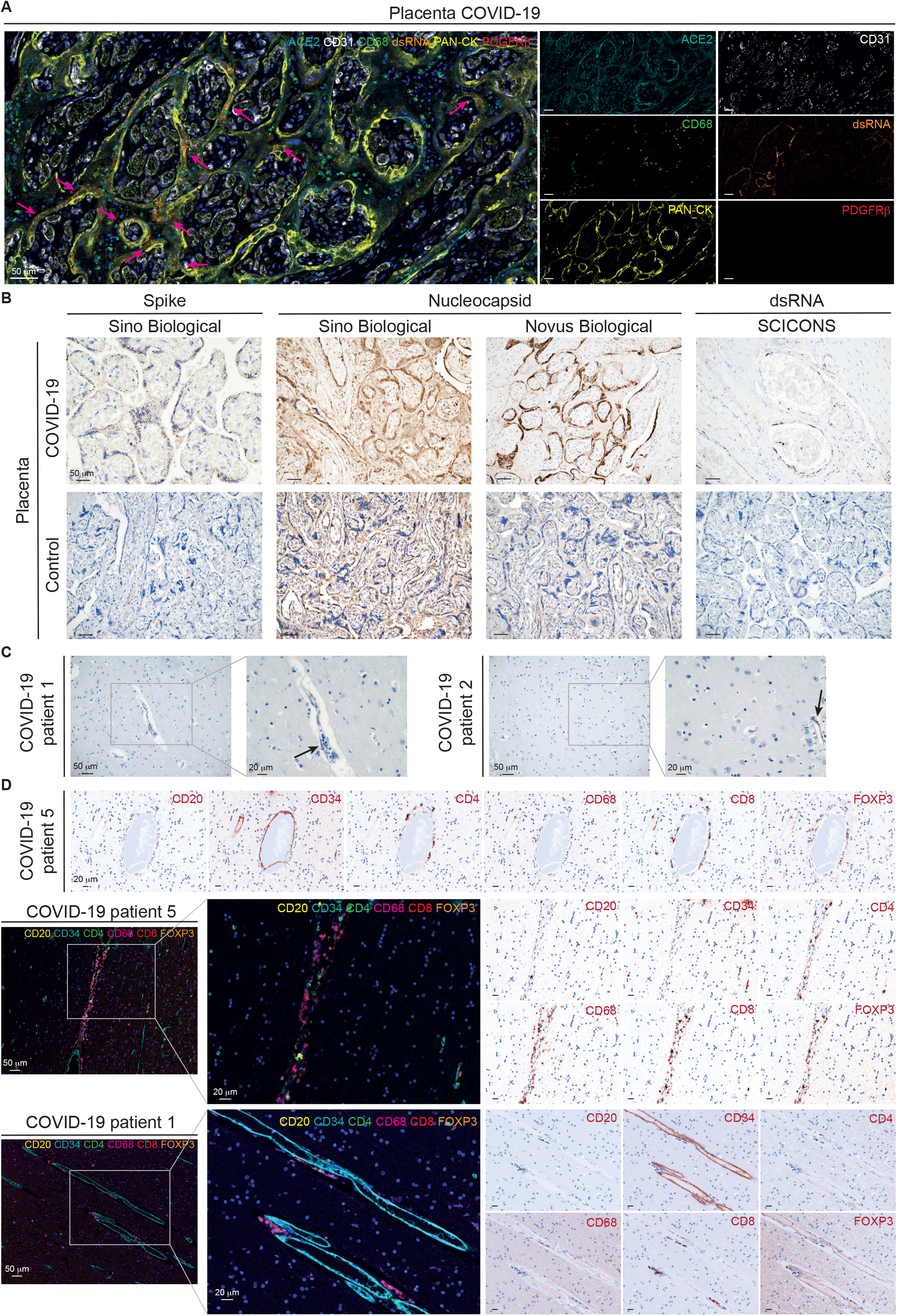
(**A**) Additional field of the 7-plex mIHC staining panel of placental tissue infected with SARS-CoV-2 presented in Figure 2D. The magenta arrows indicate accumulation of viral dsRNA in correspondence of the ACE2-positive areas by the specialized epithelial layer of syncytiotrophoblast in the placenta. The intensity of each OPAL fluorophore is further presented in individual photomicrographs. (**B**) Immunohistochemical detection of viral components and dsRNA in a COVID-19-infected placenta and in a normal placental specimen. Cell nuclei are counterstained with hematoxylin (blue). (**D**) Multiplex IHC fields of the peri-vascular immune cell infiltration in the frontal cortex of two COVID-19 patients. The antibody panel was designed for the concomitant detection of CD34 (endothelium) and five immune cell markers: CD4 (T helper cells), CD8 (cytotoxic T lymphocytes), CD20 (B cells), CD68 (macrophages), and FOXP3 (regulatory T cells). Individual OPAL intensities are displayed as “pathology view”, in which the fluorescent signal is transformed into a digitalized chromogenic DAB-like deposit.

**Supplementary Figure 3.**
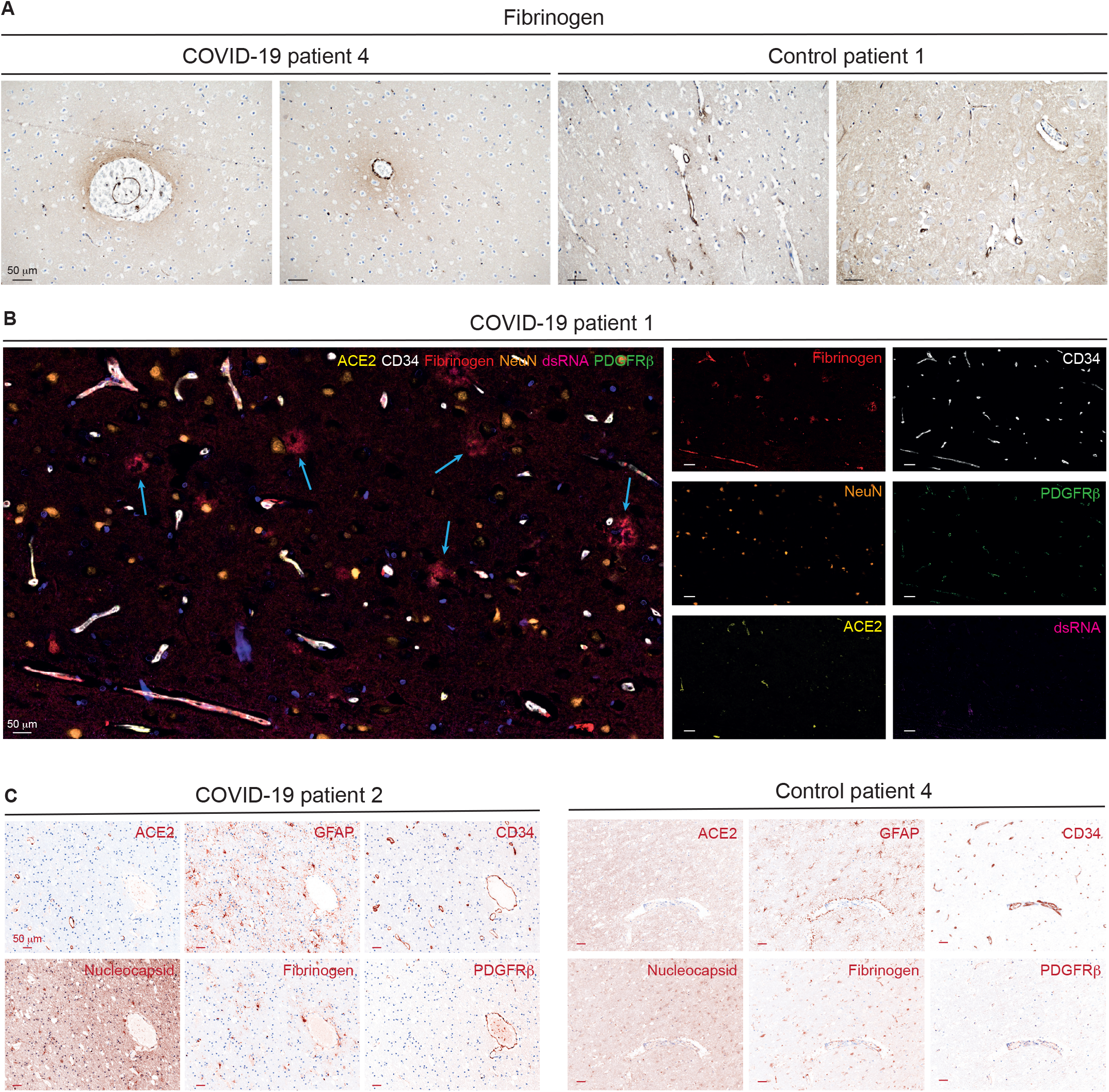
(**A**) Representative IHC of fibrinogen staining in the brain of a COVID-19 patients and in control tissue. Cell nuclei are counterstained with hematoxylin (blue). (**B**) Additional field of the composite mIHC in the COVID-19 patient presented in Figure 3A. The fields depict the neurovascular unit (CD34, PDGFRβ, and ACE2), fibrinogen, viral dsRNA, and neurons. The cyan arrows indicate fibrinogen leakage The intensity of each OPAL fluorophore is further presented in individual photomicrographs. (**C**) Individual OPAL intensities of the two panels presented in Figure 3A are displayed as “pathology view”, in which the fluorescent signal is transformed into a digitalized chromogenic DAB-like deposit.

**Supplementary Table 1.** **C**haracteristics of the **COVID-19** patients and relative controls (brain and placenta **IHC**/m**IHC** study, **L**und **U**niversity)

| Patient ID | Age (years) | Gender | ACE2 | Vascular histopathological evaluation | Neurological assessment |
| --- | --- | --- | --- | --- | --- |
| COVID-19 1 | 67 | M | High | Small vessel disease, signs of cerebral hypoperfusion | Unconscious >4 weeks before death |
| COVID-19 2 | 54 | M | High | Mild but general edema of CNS | Unconscious >4 weeks before death |
| COVID-19 3 | 72 | M | Moderate | No remarks | Unconscious >1 week before death |
| COVID-19 4 | 80 | M | Not detected | No remarks | Alert except for two days of confusion |
| COVID-19 5 | 57 | M | Not detected | Perivascular lymphocytic infiltration/vascular thrombosis | Alert until death |
| COVID-19 6 | 75 | F | Not detected | No remarks | Alert until death |
| Control 1 | 73 | M | Not detected | No remarks | No remarks, Alert until death |
| Control 2 | 68 | F | High | Hypertensive angiopathy and microthrombosis | Decline before death |
| Control 3 | 72 | F | Low | Severe vascular/ischemic lesions | No remarks |
| Control 4 | 82 | F | High | Progressive multifocal leukoencephalopathy | No remarks |
| Control 5 | 37 | M | Low | No remarks | Alert until death |
| Control 6 | 68 | F | Low | Small vessel disease, microbleeds, stroke in brainstem | No remarks |
| Control 7 | 29 | M | Low | No remarks | No remarks |
| COVID-19 placenta | 27 | F | High | Not applicable | Not applicable |
| Control placenta | 28 | F | High | Not applicable | Not applicable |

**Supplementary Table 2.** **C**haracteristics of the **COVID-19** patients and relative controls (**CSF** study, **S**ahlgrenska **U**niversity hospital, **G**othenburg)

| Patient ID | Age (years) | Gender | PDGFRβ (pg/ml) | Interval<sup>a</sup> (days) | Neurological assessment |
| --- | --- | --- | --- | --- | --- |
| COVID-19 CSF 1 | 84 | F | 1065 | 10 | Confusion |
| COVID-19 CSF 2 | 70 | M | 459 | 6 | Confusion, fatigue, delayed response to questions |
| COVID-19 CSF 3 | 62 | M | 976 | 15 | Confusion, personality changes, delayed response to questions, disorientation |
| COVID-19 CSF 4 | 64 | M | 275 | 7 | Altered consciousness/somnolence, neck stiffness, photophobia |
| COVID-19 CSF 5 | 42 | M | 602 | 12 | Exhaustion, dysgeusia |
| COVID-19 CSF 6 | 72 | M | 772 | 12 | Confusion, delayed response to questions |
| COVID-19 CSF 7 | 59 | F | 678 | 13 | Confusion, fatigue, dysarthria, myoclonus |
| COVID-19 CSF 8 | 78 | F | 611 | 10 | Confusion |
| Control CSF 1 | 72 | F | 1308 | Not applicable | Not applicable |
| Control CSF 2 | 76 | M | 1938 | Not applicable | Not applicable |
| Control CSF 3 | 56 | M | 705 | Not applicable | Not applicable |
| Control CSF 4 | 72 | M | 738 | Not applicable | Not applicable |
| Control CSF 5 | 47 | M | 1246 | Not applicable | Not applicable |
| Control CSF 6 | 73 | M | 1051 | Not applicable | Not applicable |
| Control CSF 7 | 59 | F | 905 | Not applicable | Not applicable |
| Control CSF 8 | 67 | F | 941 | Not applicable | Not applicable |

<sup>a</sup>Days between the initial report of COVID-19 symptoms and the lumbar puncture.

**Supplementary Table 3.** **A**ntibody list

| Target | Manufacturer | Article number | Host | Reactivity | Clonality | Dilution | HIER<sup>a</sup> | Application |
| --- | --- | --- | --- | --- | --- | --- | --- | --- |
| ACE2 | Abcam | Ab108252 | Rabbit | Human, Mouse, Rat | Monoclonal EPR4435[2] | 1:100 | Citrate buffer (pH6) | IHC/mlHC |
| ACE2 | R&D | MAB933 | Mouse | Human, Hamster | Monoclonal 171606 | 1:50 | Tris-EDTA buffer (pH9) | IHC |
| CD4 | Akoya Biosciences | Kit: OP7TL4001KT |  | Human |  | 70 ng/ml | Tris-EDTA buffer (pH9) | mlHC |
| CD8 | Akoya Biosciences | Kit: OP7TL4001KT |  | Human |  | 120 ng/ml | Tris-EDTA buffer (pH9) | mlHC |
| CD20 | Akoya Biosciences | Kit: OP7TL4001KT |  | Human |  | 100 ng/ml | Citrate buffer (pH6) | mlHC |
| CD31 | Cell Signaling Technology | 3528S | Mouse | Human | Monoclonal 89C2 | 1:1000 | Citrate buffer (pH6) | mlHC |
| CD34 | Novus Biologicals | NBP2-32932 | Mouse | Human, Primate | Monoclonal QBEnd/10 | 1:1000 | Citrate buffer (pH6) | mlHC |
| CD68 | Akoya Biosciences | Kit: OP7TL4001KT |  | Human |  | 15 ng/ml | Citrate buffer (pH6) | mlHC |
| dsRNA | SCICONS | 10010200 | Mouse | Virus | Monoclonal J2 | 1:500 | Tris-EDTA buffer (pH9) | IHC/mlHC |
| Fibrinogen | Abcam | Ab189490 | Rabbit | Human, Mouse, Rat | Monoclonal EPR18145-84 | 1:4000 | Tris-EDTA buffer (pH9) | IHC/mlHC |
| FOXP3 | Akoya Biosciences | Kit: OP7TL4001KT |  | Human |  | 700 ng/ml | Citrate buffer (pH6) | mlHC |
| GFAP | Atlas Antibodies | HPA056030 | Rabbit | Human, Mouse |  | 1:2500 | Citrate buffer (pH6) | IHC/mlHC |
| NeuN/RBFOX3 | Atlas Antibodies | HPA030790 | Rabbit | Human | Polyclonal | 1:200 | Citrate buffer (pH6) | mlHC |
| pan-CK | Akoya Biosciences | Kit: OP7TL4001KT |  | Human |  | 400 ng/ml | Citrate buffer (pH6) | mlHC |
| PDGFRβ | Cell Signaling Technology | 3169S | Rabbit | Human, Mouse, Rat | Monoclonal 28E1 | 1:75 | Tris-EDTA buffer (pH9) | mlHC |
| SARS-CoV nucleocapsid | Novus Biologicals | NB100-56576SS | Rabbit | Virus | Polyclonal | 1:100 | Citrate buffer (pH6) | IHC/mlHC |
| SARS-CoV-2 nucleocapsid | Sino Biologicals | 40143-019 | Rabbit | Virus | Monoclonal 019 | 1:100 | Citrate buffer (pH6) | IHC |
| SARS-CoV-2 spike | GeneTex | GTX135356 | Rabbit | Virus | Polyclonal | 1:100 | Citrate buffer (pH6) | IHC/mlHC |
| SARS-CoV-2 spike | Sino Biological | 40150-R007 | Rabbit | Virus | Monoclonal 007 | 1:100 | Citrate buffer (pH6) | IHC/mlHC |

<sup>a</sup>HIER: heat-induced epitope retrieval

**Supplementary Table 4.** **M**ultiplex **IHC** panels

| Panel | Position 1 | Position 2 | Position 3 | Position 4 | Position 5 | Position 6 |
| --- | --- | --- | --- | --- | --- | --- |
| **Neurovascular unit (Fig. 1E)** |  |  |  |  |  |  |
| HIER<sup>a</sup>/Stripping | pH9 | pH6 | pH6 |  |  |  |
| Blocking |  | 10 min, RT |  |  |  |  |
| Primary antibody | PDGFRβ | ACE2 | CD31 | | | |
| Dilution | 1:75 | 1:100 | 1:1000 |  |  |  |
| Incubation (time, temperature) |  | 30 min, RT |  |  |  |  |
| OPAL | OPAL 480 | OPAL 620 | OPAL 690 |  |  |  |
| Dilution | 1:100 | 1:100 | 1:100 |  |  |  |
| Incubation (time, temperature) |  | 10 min, RT |  |  |  |  |
| Panel | Position 1 | Position 2 | Position 3 | Position 4 | Position 5 | Position 6 |
| **Placenta (Fig. 2B)** |  |  |  |  |  |  |
| HIER/Stripping | pH9 | pH9 | pH6 | pH6 | pH6 | pH6 |
| Blocking |  |  | 10 min, RT |  |  |  |
| Primary antibody | dsRNA | PDGFRβ | ACE2 | Pan-CK | CD68 | CD31 |
| Dilution | 1:500 | 1:75 | 1:100 |  |  | 1:1000 |
| Incubation (time, temperature) |  |  | 30 min, RT |  |  |  |
| OPAL | OPAL 480 | OPAL 520 | OPAL 570 | OPAL 620 | OPAL 690 | OPAL 780 |
| Dilution | 1:100 | 1:50 | 1:50 | 1:50 | 1:50 | 1:25 |
| Incubation (time, temperature) |  |  | 10 min, RT |  |  |  |
| Panel | Position 1 | Position 2 | Position 3 | Position 4 | Position 5 | Position 6 |
| **Immunology (Fig. 2E)** |  |  |  |  |  |  |
| HIER/Stripping | pH9 | pH9 | pH6 | pH6 | pH6 | pH6 |
| Blocking |  |  | 10 min, RT |  |  |  |
| Primary antibody | CD4 | CD8 | CD20 | FOXP3 | CD68 | CD34 |
| Dilution | 70 ng/ml | 120 ng/ml | 100 ng/ml | 700 mg/ml | 15 ng/ml | 1:1000 |
| Incubation (time, temperature) |  |  | 30 min, RT |  |  |  |
| OPAL | OPAL 520 | OPAL 570 | OPAL 540 | OPAL 620 | OPAL 650 | OPAL 690 |
| Dilution | 1:50 | 1:50 | 1:50 | 1:50 | 1:50 | 1:50 |
| Incubation (time, temperature) |  |  | 10 min, RT |  |  |  |
| Panel | Position 1 | Position 2 | Position 3 | Position 4 | Position 5 | Position 6 |
| **Fibrinogen (Fig. 3A)** |  |  |  |  |  |  |
| HIER/Stripping | pH9 | pH9 | pH9 | pH6 | pH6 | pH6 |
| Blocking |  |  | 10 min, RT |  |  |  |
| Primary antibody | dsRNA | PDGFRβ | Fibrinogen | ACE2 | NeuN | CD34 |
| Dilution | 1:500 | 1:75 | 1:4000 | 1:100 | 1:200 | 1:1000 |
| Incubation (time, temperature) |  |  | 30 min, RT |  |  |  |
| OPAL | OPAL 480 | OPAL 520 | OPAL 570 | OPAL 620 | OPAL 690 | OPAL 780 |
| Dilution | 1:100 | 1:50 | 1:50 | 1:50 | 1:50 | 1:25 |
| Incubation (time, temperature) |  |  | 10 min, RT |  |  |  |
| Panel | Position 1 | Position 2 | Position 3 | Position 4 | Position 5 | Position 6 |
| **Astrocytes (Fig. 3B)** |  |  |  |  |  |  |
| HIER/Stripping | pH9 | pH9 | pH6 | pH6 | pH6 | pH6 |
| Blocking |  |  | 10 min, RT |  |  |  |
| Primary antibody | PDGFRβ | Fibrinogen | ACE2 | GFAP | CoV-2 spike | CD34 |
| Dilution | 1:75 | 1:4000 | 1:100 | 1:2500 | 1:100 | 1:1000 |
| Incubation (time, temperature) |  |  | 30 min, RT |  |  |  |
| OPAL | OPAL 520 | OPAL 570 | OPAL 540 | OPAL 620 | OPAL 650 | OPAL 690 |
| Dilution | 1:50 | 1:50 | 1:50 | 1:50 | 1:50 | 1:50 |
| Incubation (time, temperature) |  |  | 10 min, RT |  |  |  |

<sup>a</sup>HIER: heat-induced epitope retrieval

## References

1. Chen N, Zhou M, Dong X, et al. Epidemiological and clinical characteristics of 99 cases of 2019 novel coronavirus pneumonia in Wuhan, China: a descriptive study. Lancet. Feb 15 2020;395(10223):507–513. doi:110.1016/S0140-6736(20)30211-7

2. Wang D, Hu B, Hu C, et al. Clinical Characteristics of 138 Hospitalized Patients With 2019 Novel Coronavirus-Infected Pneumonia in Wuhan, China. JAMA. Mar 17 2020;323(11):1061–1069. doi:10.1001/jama.2020.1585

3. Puelles VG, Lutgehetmann M, Lindenmeyer MT, et al. Multiorgan and Renal Tropism of SARS-CoV-2. N Engl J Med. Aug 6 2020;383(6):590–592. doi:10.1056/NEJMc2011400

4. Gupta A, Madhavan MV, Sehgal K, et al. Extrapulmonary manifestations of COVID-19. Nat Med. Jul 2020;26(7):1017–1032. doi:10.1038/s41591-020-0968-3

5. Paterson RW, Brown RL, Benjamin L, et al. The emerging spectrum of COVID-19 neurology: clinical, radiological and laboratory findings. Brain. Oct 1 2020;143(10):3104–3120. doi:10.1093/brain/awaa240

6. Helms J, Kremer S, Merdji H, et al. Neurologic Features in Severe SARS-CoV-2 Infection. N Engl J Med. Jun 4 2020;382(23):2268–2270. doi:10.1056/NEJMc2008597

7. Mao L, Jin H, Wang M, et al. Neurologic Manifestations of Hospitalized Patients With Coronavirus Disease 2019 in Wuhan, China. JAMA Neurol. Jun 1 2020;77(6):683–690. doi:10.1001/jamaneurol.2020.1127

8. Oxley TJ, Mocco J, Majidi S, et al. Large-Vessel Stroke as a Presenting Feature of Covid-19 in the Young. N Engl J Med. May 14 2020;382(20):e60. doi:10.1056/NEJMc2009787

9. Hoffmann M, Kleine-Weber H, Schroeder S, et al. SARS-CoV-2 Cell Entry Depends on ACE2 and TMPRSS2 and Is Blocked by a Clinically Proven Protease Inhibitor. Cell. Apr 16 2020;181(2):271–280 e8. doi:10.1016/j.cell.2020.02.052

10. Li MY, Li L, Zhang Y, Wang XS. Expression of the SARS-CoV-2 cell receptor gene ACE2 in a wide variety of human tissues. Infect Dis Poverty. Apr 28 2020;9(1):45. doi:10.1186/s40249-020-00662-x

11. Tabula Muris C, Overall c, Logistical c, et al. Single-cell transcriptomics of 20 mouse organs creates a Tabula Muris. Nature. Oct 2018;562(7727):367–372. doi:10.1038/s41586-018-0590-4

12. Vanlandewijck M, He L, Mae MA, et al. A molecular atlas of cell types and zonation in the brain vasculature. Nature. Feb 22 2018;554(7693):475–480. doi:10.1038/nature25739

13. He L, Vanlandewijck M, Mae MA, et al. Single-cell RNA sequencing of mouse brain and lung vascular and vessel-associated cell types. Sci Data. Aug 21 2018;5:180160. doi:10.1038/sdata.2018.160

14. Lein ES, Hawrylycz MJ, Ao N, et al. Genome-wide atlas of gene expression in the adult mouse brain. Nature. Jan 11 2007;445(7124):168–76. doi:10.1038/nature05453

15. Allen Mouse Brain Atlas [Internet]. Seattle (WA): Allen Institute for Brain Science. 2011. Available from: .https://portal.brain-map.org/atlases-and-data/rnaseq/human-multiple-cortical-areas-smart-seq

16. Uhlen M, Fagerberg L, Hallstrom BM, et al. Proteomics. Tissue-based map of the human proteome. Science. Jan 23 2015;347(6220):1260419. doi:10.1126/science.1260419

17. Miners JS, Kehoe PG, Love S, Zetterberg H, Blennow K. CSF evidence of pericyte damage in Alzheimer’s disease is associated with markers of blood-brain barrier dysfunction and disease pathology. Alzheimers Res Ther. Sep 14 2019;11(1):81. doi:10.1186/s13195-019-0534-8

18. Doobay MF, Talman LS, Obr TD, Tian X, Davisson RL, Lazartigues E. Differential expression of neuronal ACE2 in transgenic mice with overexpression of the brain reninngiotensin system. Am J Physiol Regul Integr Comp Physiol. Jan 2007;292(1):R373–81. doi:10.1152/ajpregu.00292.2006

19. Xu J, Lazartigues E. Expression of ACE2 in Human Neurons Supports the Neuro-Invasive Potential of COVID-19 Virus. Cell Mol Neurobiol. Jul 4 2020;doi:10.1007/s10571-020-00915-1

20. Gallagher PE, Chappell MC, Ferrario CM, Tallant EA. Distinct roles for ANG II and ANG-(1-7) in the regulation of angiotensin-converting enzyme 2 in rat astrocytes. Am J Physiol Cell Physiol. Feb 2006;290(2):C420–6. doi:10.1152/ajpcell.00409.2004

21. Hamming I, Timens W, Bulthuis ML, Lely AT, Navis G, van Goor H. Tissue distribution of ACE2 protein, the functional receptor for SARS coronavirus. A first step in understanding SARS pathogenesis. J Pathol. Jun 2004;203(2):631–7. doi:10.1002/path.1570

22. McCracken IR, Saginc G, He L, et al. Lack of Evidence of ACE2 Expression and Replicative Infection by SARSCoV-2 in Human Endothelial Cells. Circulation. Jan 6 2021;doi:10.1161/CIRCULATIONAHA.120.052824

23. Matschke J, Lutgehetmann M, Hagel C, et al. Neuropathology of patients with COVID-19 in Germany: a post-mortem case series. Lancet Neurol. Nov 2020;19(11):919–929. doi:10.1016/S1474-4422(20)30308-2

24. Solomon IH, Normandin E, Bhattacharyya S, et al. Neuropathological Features of Covid-19. N Engl J Med. Sep 3 2020;383(10):989–992. doi:10.1056/NEJMc2019373

25. Bradley BT, Maioli H, Johnston R, et al. Histopathology and ultrastructural findings of fatal COVID-19 infections in Washington State: a case series. Lancet. Aug 1 2020;396(10247):320–332. doi:10.1016/S0140-6736(20)31305-2

26. Zaigham M, Holmberg A, Karlberg ML, et al. Intrauterine vertical SARS-CoV-2 infection: a case confirming transplacental transmission followed by divergence of the viral genome. BJOG. Feb 27 2021;doi:10.1111/1471-0528.16682

27. Lee M-H, Perl DP, Nair G, et al. Microvascular Injury in the Brains of Patients with Covid-19. New England Journal of Medicine. 2020;doi:10.1056/NEJMc2033369

28. Thakur KT, Miller EH, Glendinning MD, et al. COVID-19 neuropathology at Columbia University Irving Medical Center/New York Presbyterian Hospital. Brain. Apr 15 2021;doi:10.1093/brain/awab148

29. Pilotto A, Masciocchi S, Volonghi I, et al. SARS-CoV-2 encephalitis is a cytokine release syndrome: evidences from cerebrospinal fluid analyses. Clin Infect Dis. Jan 4 2021;doi:10.1093/cid/ciaa1933

30. Eden A, Kanberg N, Gostner J, et al. CSF biomarkers in patients with COVID-19 and neurological symptoms: A case series. Neurology. Oct 1 2020;doi:10.1212/WNL.0000000000010977

31. Banerjee AK, Blanco MR, Bruce EA, et al. SARS-CoV-2 Disrupts Splicing, Translation, and Protein Trafficking to Suppress Host Defenses. Cell. Nov 25 2020;183(5):1325–1339 e21. doi:10.1016/j.cell.2020.10.004

32. Rhea EM, Logsdon AF, Hansen KM, et al. The S1 protein of SARS-CoV-2 crosses the blood-brain barrier in mice. Nat Neurosci. Dec 16 2020;doi:10.1038/s41593-020-00771-8

33. Armulik A, Genove G, Mae M, et al. Pericytes regulate the blood-brain barrier. Nature. Nov 25 2010;468(7323):557–61. doi:10.1038/nature09522

34. Wahlgren N, Thoren M, Hojeberg B, et al. Randomized assessment of imatinib in patients with acute ischaemic stroke treated with intravenous thrombolysis. J Intern Med. Mar 2017;281(3):273–283. doi:10.1111/joim.12576

35. Su EJ, Fredriksson L, Geyer M, et al. Activation of PDGF-CC by tissue plasminogen activator impairs blood-brain barrier integrity during ischemic stroke. Nat Med. Jul 2008;14(7):731–7. doi:10.1038/nm1787

